# Self-induced Dimensional Reduction and Scaling Transition of mRNA in Polysomes: A Multiscale Simulation Study

**DOI:** 10.64898/2026.01.26.701782

**Authors:** Hideki Kobayashi, Horacio V. Guzman

**Affiliations:** Institute for Computational Physics (ICP), University of Stuttgart, Allmandring 3, 70569 Stuttgart, Germany; Institut de Ciència de Materials de Barcelona ICMAB-CSIC, Campus de la UAB, E-08193 Spain

## Abstract

The spatial architecture and mechanical rigidity of polysomes are crucial determinants of translational efficiency and mRNA stability. In this study, we investigate the conformational statistics of an mRNA backbone decorated with high-density ribosomes at varying densities using large-scale, extensive molecular dynamics simulations based on the Kremer-Grest bead-spring model. To address the extreme spatial asymmetry between mRNA monomers and ribosomes, we used an efficient tree-based neighbour list algorithm, enabling the analysis of mRNA chains up to *N* = 4, 969. Our results demonstrate that the excluded volume of massive ribosomes induces a significant and robust expansion of the scaling exponent *v* from 0.59 to approximately 0.7. In the conformation of mRNA, this shift translates to a self-induced dimensional reduction from a three-dimensional random coil toward a stretched, a quasi-two-dimensional architecture at biologically relevant scales. Such a transition is further evidenced by a periodic “regain” of the bond-bond correlation function *C*(*n*) at ribosome attachment sites, indicating a geometric alignment absent in standard homopolymers. These findings reveal that the geometric crowding of ribosomes itself provides a robust physical prerequisite for the formation of higher-order polysome architectures, bridging the gap between polymer physics and structural properties of mRNA during translation.

## 1. INTRODUCTION

The spatial architecture and mechanical rigidity of polysomes, which are complexes formed by multiple ribosomes translating a single mRNA strand, are fundamental determinants of protein synthesis regulation and mRNA stability. Recent cryo-electron tomography studies have revealed that polysomes can adopt sophisticated higher-order structures, such as helical or circular configurations, including the notable “closed-loop” model, among others1–8. These structures are thought to optimise translation efficiency and protect genetic information from enzymatic degradation^9–13^. From a physical perspective, the maintenance of these architectures depends critically on the effective stiffness of the mRNA backbone. However, the microscopic origin of this rigidity remains poorly understood, often being reduced to a generic increase in persistence length without a rigorous statistical-mechanical basis.

Ribosome attachment is expected to influence mRNA conformation through two distinct physical mechanisms: (i) the enhancement of local bending rigidity due to the binding of massive protein complexes, and (ii) the global steric exclusion imposed by the large ribosomal volumes. Previous numerical studies, such as those by Fernandes et al.^14^, have focused on the former, premising that ribosomal attachment increases the persistence length and thereby determines the mRNA’s spatial extent. To isolate these effects, it is helpful to consider a “ghost” ribosome model as a baseline, where appendages increase local stiffness at their attachment points but lack mutual steric repulsion, although in actual cellular environments, ribosomes are bulky structures that are significantly larger than the mRNA strand.

This observation raises a fundamental question we tackle: how does the ribosomal excluded volume affect mRNA during translation? In classical polymer physics, theoretical and experimental work on bottle-brush polymers suggests that flexible side chains merely rescale the backbone’s persistence length without altering its scaling exponents^15–17^; any observed shift in *ν* is typically attributed to finite-size effects rather than a change in the universality class. However, ribosomes, exceeding the mRNA monomer diameter by more than thirty-fold, are not only massive but are also, comparatively, highly rigid macromolecules. We argue that the interplay of extreme asymmetry and rigidity creates collective multi-body interactions among ribosomes that can no longer be mapped onto a single stiffness parameter. These collective interactions construct a “steric corridor” that non-locally restricts the backbone conformational space, which could shift the polymer into a different universality class. Conventional models that neglect these interactions risk failing to capture the collective behaviour of the polysome as a supramolecular assembly. Nevertheless, this effect remains unexplored because simulating such extreme asymmetries in the long-chain limit is computationally challenging and demands new modeling schemes.

In this work, we investigate the conformational statistics of mRNA in polysomes by employing the standard Kremer-Grest (KG) bead-spring model^18^. By combining the Weeks-Chandler-Andersen (WCA) and FENE potentials, we strip away chemical specificity to isolate the pure role of geometric constraints. Our focus is on how the spatial asymmetry between mRNA monomers and ribosomes dictates the global scaling behavior. However, we find that the standard cell-list algorithm faces significant practical challenges in such a spatially asymmetric system. Conventional grid-based approaches often encounter computational and memory bottlenecks when attempting to simulate full-length mRNA (*N* = 4, 969) while simultaneously handling a large number of polysomes to ensure statistical robustness.

Depending on the hardware environment, these overheads can make it difficult to achieve the necessary sampling within a reasonable timeframe. To overcome these constraints, we use an efficient tree-based neighbor-list algorithm. Unlike the standard method, the tree-based approach adaptively manages the disparate length scales across the polysome, enabling precise and rapid evaluation of steric repulsions. This technical advancement facilitates the large-scale simulations required to maintain the low-density limit, thereby ensuring that our scaling analysis remains statistically independent and robust.

Our results demonstrate that the transition of the mRNA backbone toward a stretched chain regime (*v*≈ 0.7) is not merely a consequence of local stiffening, but is fundamentally driven by a “self-induced dimensional reduction” caused by the crowded arrangement of real ribosomal volumes. This geometric alignment serves as a robust foundation for the formation of complex polysome architectures.

## II. MODEL AND NUMERICAL METHOD

In this study, we construct a coarse-grained model to simulate the conformational behaviour of a polysome (see Fig. 1A), defined as an mRNA molecule associated with multiple ribosomes. This model aims to capture essential features of polysome organisation and the spatial configuration of mRNA under translationally active conditions.

**FIG. 1.**
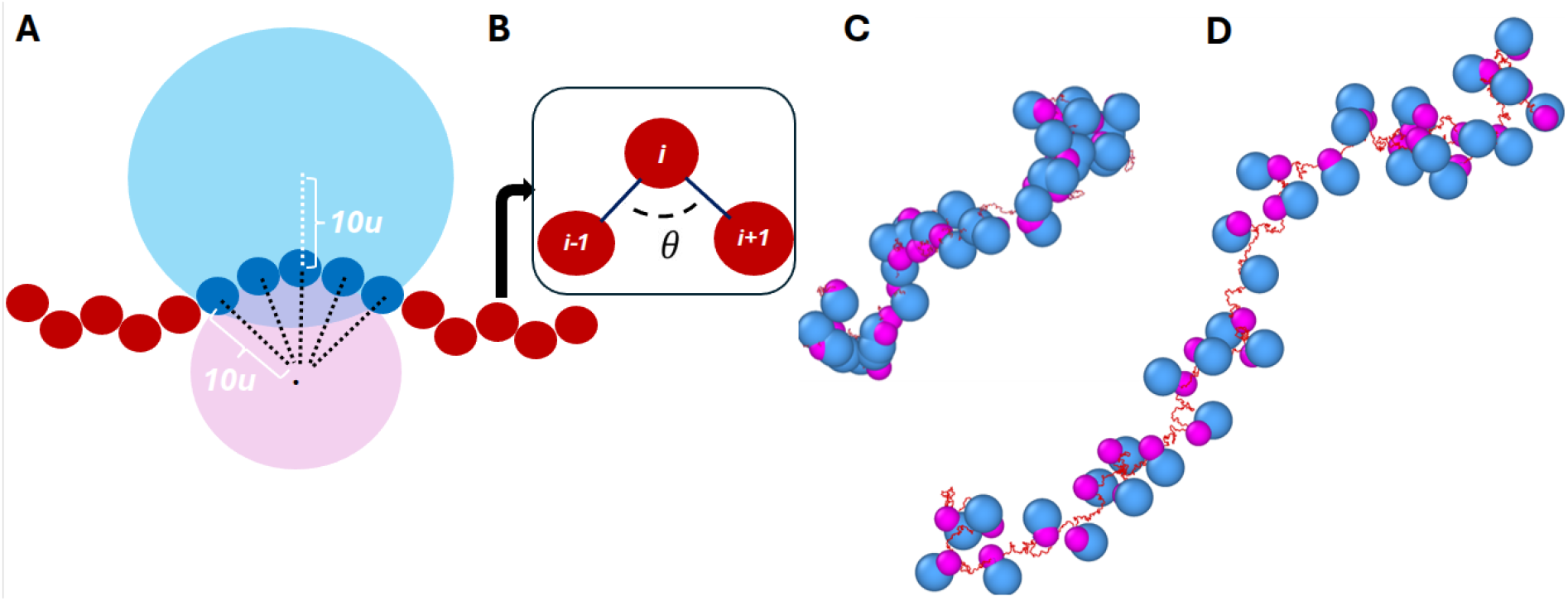
(A) Schematic illustration of the coarse-grained ribosome-mRNA model. The ribosome consists of a large subunit (cyan) and a small subunit (pink). The mRNA backbone is represented by a chain of monomers (red). Within the 30-monomer footprint, monomers (blue) are harmonically constrained to the center of the small subunit (black dashed lines) to simulate surface attachment. For clarity, only 5 monomers are illustrated to represent the 30-monomer segment. Additionally, the central monomer of this segment is anchored to the center of the large subunit via a harmonic potential (white dotted line). This dual-anchoring construction ensures that the mRNA is sandwiched between the subunits, effectively forming a single functional unit with an elongated geometry. (B) Schematic illustration of the bending angle definition. Three consecutive monomers, indexed as *i*− 1, *i*, and *i* + 1, define the local configuration of the mRNA backbone. The bending angle *θ* is the angle formed by the two bond vectors,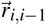 and 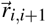 originating from monomer *i*. (C) and (D), show equilibrated mRNA-ribosome conformations. Both snapshots show a system with *N* = 4, 969, *N*_space_ = 5, and *θ* = 2*π*/3. (C) In the ghost-ribosome model, the lack of inter-ribosomal excluded volume leads to a compact mRNA configuration. (D) In the real-ribosome case, repulsive interactions induce a structural expansion, resulting in the observed non-SAW scaling behavior.

The mRNA is modelled as a polymer chain using the Kremer–Grest model^18^. Each monomer represents a coarse-grained nucleotide unit and is connected to its neighbours by finitely extensible nonlinear elastic (FENE) bonds:

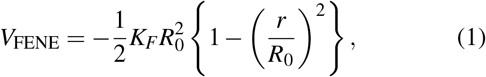

where *K*_*F*_ and *R*_0_ stand for the spring constant and the maximum extension of the bond, respectively. Excluded volume interactions between monomers are introduced via the Weeks-Chandler-Andersen potential (WCA):

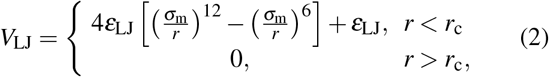

The parameters *ε*_LJ_ and *σ*_m_ are the strength of the interaction and the diameter of the monomers, respectively. *r*_c_ = 2^1/6^*σ*_m_ stands for the cutoff length of potential. The total number of monomers in the mRNA chain is represented by *N*. The simulations are performed with the following chain lengths: *N* = 2000, 3000, 4060, 4969. The diameter of each monomer is defined as the unit length *u* = 0.6*nm* in real space, and all physical quantities are expressed in reduced units based on this length. The energy unit is *k*_*B*_*T*, and the mass unit corresponds to the mass of a monomer.

We use a bending-angle potential to introduce chain stiffness and control mRNA flexibility. This potential acts on every triplet of consecutive monomers as shown in Fig. 1B, imposing a preferred bending angle *θ*_0_ using a harmonic angle potential:

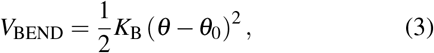

In our simulations, we consider *θ*_0_ = *π*/2 and 2*π*/3 to mimic various structural constraints. The chosen angles are based on a previous work^19^ where different mRNAs where the end-to-end distances were measured, and a simple polymeric model was used to identify the angular restrictions that can recover experimental lengths.

Each ribosome is modeled as a pair of spheres representing the small and large subunits, with diameter *σ*_S_ = 20*u*, and *σ*_L_ = 30*u* (i.e., 12 *nm* and 18 *nm*). When these two spheres are in contact, the resulting composite structure spans a long-axis length of 30 nm in real space. This value is about 20− 30% larger than experimentally observed bacterial ribosome sizes (20− 25, *nm*). However, this discrepancy is compensated for by a later correction described below.

As illustrated in Fig. 1A, the structural integrity of the ribosome-mRNA complex is maintained through a series of harmonic constraints. Each ribosome is associated with a fixed mRNA footprint consisting of 30 monomers. These monomers (represented by blue beads in the Fig. 1A) are tethered to the center of the small ribosomal subunit (pink) using a harmonic potential of the form:

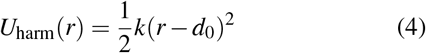

where *k* = 10 is the spring constant and *d*_0_ = 10*u* is the equilibrium distance. This potential effectively constrains the 30-monomer segment to the surface of the small subunit, simulating the physical attachment during translation.

To ensure that the two subunits behave as a single functional entity, an additional anchor is applied to the central monomer of the 30-bead footprint. This specific monomer is connected to the center of the large ribosomal subunit (cyan) via the same harmonic potential (*k* = 10, *d*_0_ = 10*u*). This dual-coupling ensures that the mRNA is effectively “sandwiched” between the subunits.

Due to this extended connection architecture, the effective long-axis length of the ribosome in our model is approximately 45*u* (27 nm), while the short-axis length remains 18 nm. Although these dimensions are slightly larger than experimental ribosome values, the short-axis alignment matches typical bacterial dimensions, providing a reasonable coarse-grained representation of the steric constraints within a polysome.

To elucidate how ribosome density dictates the spatial organization of the complex, multiple ribosomes are localized on the mRNA with a controlled separation of 30 ×*N*_space_. This allows for a detailed analysis of the interplay between ribosome spacing, RNA conformation, and the resulting polysome geometry.

All large particles (ribosomal subunits) interact via a Gaussian Core (GCP), which provides soft, long-ranged repulsion, defined as follows:

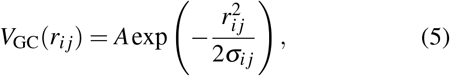

where *A* is the strength of the potential which is set to 5 for ribosome–ribosome interactions and 10 for ribosome–monomer interactions. The choice of a soft repulsive potential over a hard-core potential (e.g., WCA) is motivated by two factors. First, from a physical perspective, the interaction strength *A* = 10 provides a sufficiently high energy barrier to prevent significant overlap under physiological densities, effectively maintaining the “steric corridor” described in the Introduction. Second, from a computational standpoint, the GCP avoids the numerical instabilities and unphysical large forces that often arise in highly asymmetric systems (30:1 size ratio) when a monomer occasionally penetrates a hard-core shell. The potential is truncated and shifted to smoothly vanish at a cutoff radius set to 120% of the sum of the interacting particles’ radii *σ*_*i j*_ defined as:

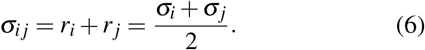

To ensure that the Gaussian potential vanishes smoothly at the cutoff radius *r*_c_, we introduce a smoothing function *S*(*r*_*i j*_), which continuously reduces the interaction strength to zero as *r*_*i j*_ → *r*_c_. The modified Gaussian Core Potential is defined as:

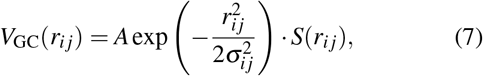

where *S*(*r*_*i j*_) is a smoothing function that satisfies *S*(*r*_*i j*_) = 1 for *r*_*i j*_ ≪*r*_*c*_ and decays smoothly to zero as *r*_*i j*_→ *r*_*c*_. We used the form of the smoothing function defined as:

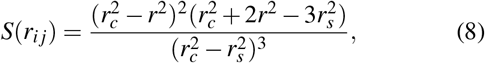

where *r*_*s*_ = 1.1× (*r*_*i*_ + *r* _*j*_) is the start of the smoothing region, and *r*_c_ is the cutoff radius, set to 1.2 × (*r*_*i*_ + *r* _*j*_). This approach ensures a smooth truncation of the potential and its derivative, improving the numerical stability of simulations. A key feature of this system is the coexistence of interaction scales, with ribosome-ribosome Gaussian potentials extending up to *r*_*c*_≈ 36*u*. While the standard cell-list method is slightly more efficient (approximately 5%) for small systems with 8 polysomes, it faces severe memory bottlenecks as the system scales. To overcome this, we employ a hierarchical tree-based neighbor list^20,21^ implemented in HOOMD-blue^22^. These performance characteristics are based on our comprehensive benchmarks conducted on the MareNostrum 5 supercomputer (using NVIDIA H100 GPUs). To evaluate scalability and optimize performance, we tested the number of polysomes from 8 up to 128 in the system and explored neighbor-list buffer distances ranging from 0.1 to 6.0*u*. We found that even in this high-performance environment, the cell-list method is restricted to a maximum of 8 polysomes due to prohibitive memory demands. In contrast, the tree-based approach enables a 2.6-fold (260%) increase in speed per-polysome calculation time and is capable of handling even larger systems. For our production runs, we chose to simulate 32 polysomes to achieve an optimal balance between computational throughput and robust statistical sampling, as the speed-up per particle starts to saturate beyond this point.

The tree-based method becomes increasingly indispensable as the monomer number *N* increases. Larger *N* not only leads to a greater end-to-end distance *R*_*E*_ but also extends the equilibration time, thereby increasing the center-of-mass excursion and the required simulation box size. To maintain the low-density limit and ensure statistical independence, each polysome is allocated a 2000^3^ *u*^3^ box. At our maximum length (*N* = 4, 969), the mean-square end-to-end distance is 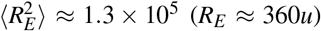 and the center-of-mass excursion reaches≈ 86*u* during equilibration. Assuming Gaussian statistics for these conformational and diffusive fluctuations, a separation of several times the standard deviation is necessary to suppress stochastic contacts between polysomes. Given that the effective interaction range can extend beyond 1000*u* under these variances, our 2000*u* box provides a robust safety margin, ensuring that the observed scaling *v* ≈0.7 is a purely intra-molecular effect. Since both neighbor-list algorithms are implemented within the HOOMD-blue framework, we have verified that they yield identical forces and energies within numerical precision.

We set the parameters *K*_F_ = 30*k*_B_*T, R*_0_ = 1.5*u, ε*_LJ_ = 1.0*k*_B_*T, K*_B_ = 50.0*k*_B_*T* and *dt* = 0.005*τ*. Using the model described above, we evaluate its structural properties via NVT molecular dynamics (MD) simulation with a Langevin thermostat at *k*_B_*T* = 1 and friction coefficient *Γ* = 0.1. This modelling framework allows us to systematically investigate how periodic ribosome binding affects mRNA conformation under various geometric constraints. The use of soft repulsive potentials enables the simulation of crowded polysomes-like conditions without hard-core overlap, while harmonic constraints impose realistic tethering of RNA to ribosomal subunits. The angle potential allows us to tune RNA stiffness and assess its impact on large-scale structure formation. Altogether, this modelling framework provides a minimal yet physically informed platform for systematically exploring RNA organisation in the context of active translation.

## III. RESULTS

To ensure that the systems are properly equilibrated, we estimate the characteristic relaxation time of a system. Specifically, we calculated the time-displaced autocorrelation function of the end-to-end vector, ⟨**R**_E_(*t*). **R**_E_(0) ⟩, for *N* = 1000 and *N*_*space*_ = 5, where **R**_E_ = **r**_*N*_− **r**_1_. As shown in Fig. 2, this function can be well approximated by an exponential decay,

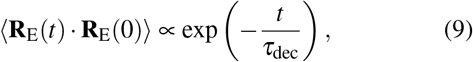

where the data indicate *τ*_dec_≈ 48000*τ* = 0.48G*N*^2^*τ*. This corresponds to approximately 15 *τ*_*R*_, where *τ*_*R*_ is the Rouse relaxation time of a corresponding flexible chain ^23^, i.e. naked RNA. While the attachment of ribosomes significantly slows down the conformational dynamics, we conclude that equilibration at least over a time *τ*_eq_ = 3*τ*_dec_ is sufficient (note exp(− 3) ≈0.05). Following this equilibration, production runs were conducted for 3.6× 10^6^*τ* (≈25*τ*_eq_) for *N* = 1000, 3.0 ×10^6^*τ* (≈16*τ*_eq_) for *N* = 2000, and 2.5 ×10^6^*τ* (≈ 5*τ*_eq_) for *N* = 3000. For the largest systems (*N* = 4060 and 4969), production runs were performed for *τ*_eq_ to balance the heavy computational cost on the supercomputer with the need for high-resolution structural analysis. To obtain reliable ensemble averages, we utilized 100 independent replicas for the naked mRNA system. For the real-ribosome system, where the complex interplay of excluded volume effects governs the scaling behavior, we ensured robust statistics by simulating 32 independent replicas. In contrast, for the ghost-ribosome system, 8 replicas were employed; as this system primarily serves as a reference to confirm the well-established theoretical prediction that side-chain presence merely renormalizes the persistence length without altering the universal scaling (*v*≈ 0.6), this sampling level was deemed sufficient to capture the qualitative trend without diverting excessive computational resources from the primary analysis of real-ribosome effects. The naked mRNA systems were simulated using ESPResSo++^24^, as its sparse interaction table efficiently manages 100 distinct particle types. In contrast, for ribosome-containing systems, we migrated to HOOMD-blue (v5) to resolve the aforementioned cell-based neighbor list bottleneck, leveraging its tree-based search instead.

**FIG. 2.**
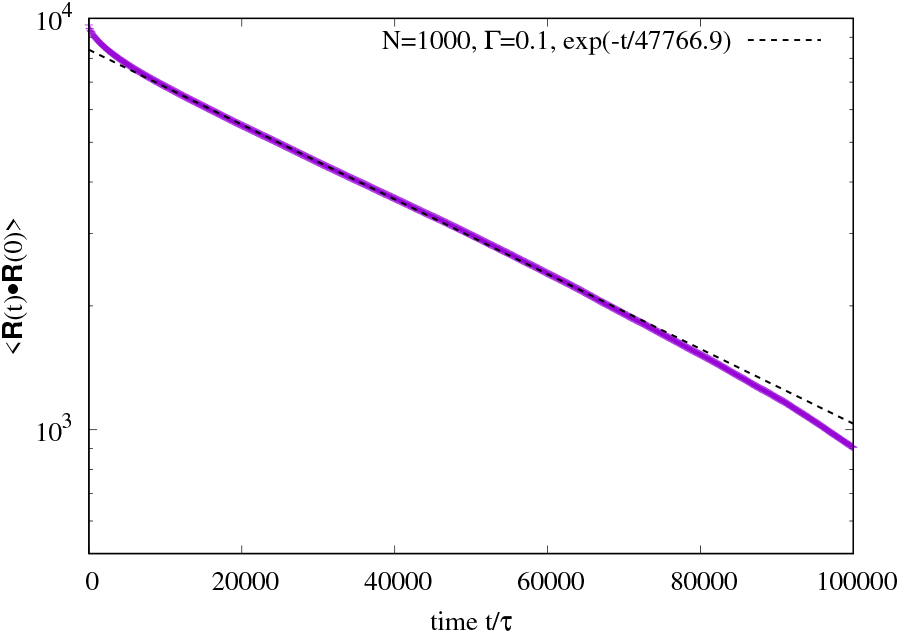
End-to-end vector autocorrelation function ⟨**R**_E_(*t*) · **R**_E_(0) ⟩ for *N* = 1000 (solid line). It is proportional to exp[− *t*/(47766.9*τ*)] (dashed line).

Before quantifying the spatial statistics, we examine the representative equilibrated conformations of the polysome. Figure 1C and D compares the mRNA-ribosome assemblies for *N* = 4, 969, *N*_space_ = 5, and *θ* = 2*π*/3 in the ghost and real ribosome cases. In the ghost ribosome system (Fig. 1(Left)), where excluded volume interactions between ribosomes are absent, the subunits frequently overlap, and the mRNA backbone adopts a compact, disordered configuration similar to a typical random coil. In contrast, the real ribosome system (Fig. 1(Right)) clearly demonstrates the impact of steric repulsion. The Gaussian core interactions prevent subunit overlapping, effectively organizing the ribosomes into a more extended chain. This steric repulsion “pushes” the intervening mRNA segments outward, driving the entire complex toward a significantly more expanded state. This visual observation provides a direct physical basis for the increased scaling exponent *v* discussed in the following section.

The scaling exponent *v* characterizes the spatial extent of a polymer chain, describing how its size (e.g., the end-to-end distance *R*_E_) scales with the number of monomers *N* as 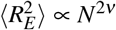. In polymer physics, *v* = 1/2 corresponds to an ideal, Gaussian chain where excluded-volume interactions are neglected. A value of *v* = 0.588 (or 3/5 in Flory theory) indicates a ‘swollen’ chain in a good solvent, where monomers repel each other. At the upper limit, *v* = 1 represents a perfectly rigid, rod-like conformation. Thus, *v* serves as a sensitive probe for the effective stiffness and the steric environment of the mRNA within the polysome.

To determine the scaling exponent *ν*, we first examine the mean-square end-to-end distance 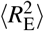 for two baseline cases: the naked mRNA and the mRNA associated with “ghost” ribosomes. In the ghost-ribosome model, the monomers are constrained to the ribosomal surface as described in the Methods, but all repulsive interactions involving the ribosomal subunits are switched off. As shown in Fig. 3(Left), the naked mRNA—representing the standard KG model with a bending potential—yields *ν* ≈ 0.6. This is consistent with the expected scaling for a self-avoiding walk in a good solvent and agrees with previous studies employing the KG model with angular constraints^25^. For the mRNA with ghost ribosomes, the scaling exponent remains *ν* ≈ 0.6, indicating that the local structural constraint imposed by the ribosome binding sites does not fundamentally affect the fractal dimension of the polymer at large length scales. However, it should be noted that the absolute magnitude of 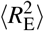 for the ghost-ribosome system is larger than that for the naked mRNA at any given N. This chain expansion can be attributed to the local stiffening and the spatial occupancy of the binding segments, which effectively increase the Kuhn length of the chain. These baseline results can be well understood through the blob picture^26^, where the ghost ribosomes act as local perturbations that renormalise the persistence length without shifting the global universality class of the self-avoiding-walk (SAW). Similar results and analysis apply for Fig. 3(Right), where the bending angle is *θ* = 2*π*/3.

**FIG. 3.**
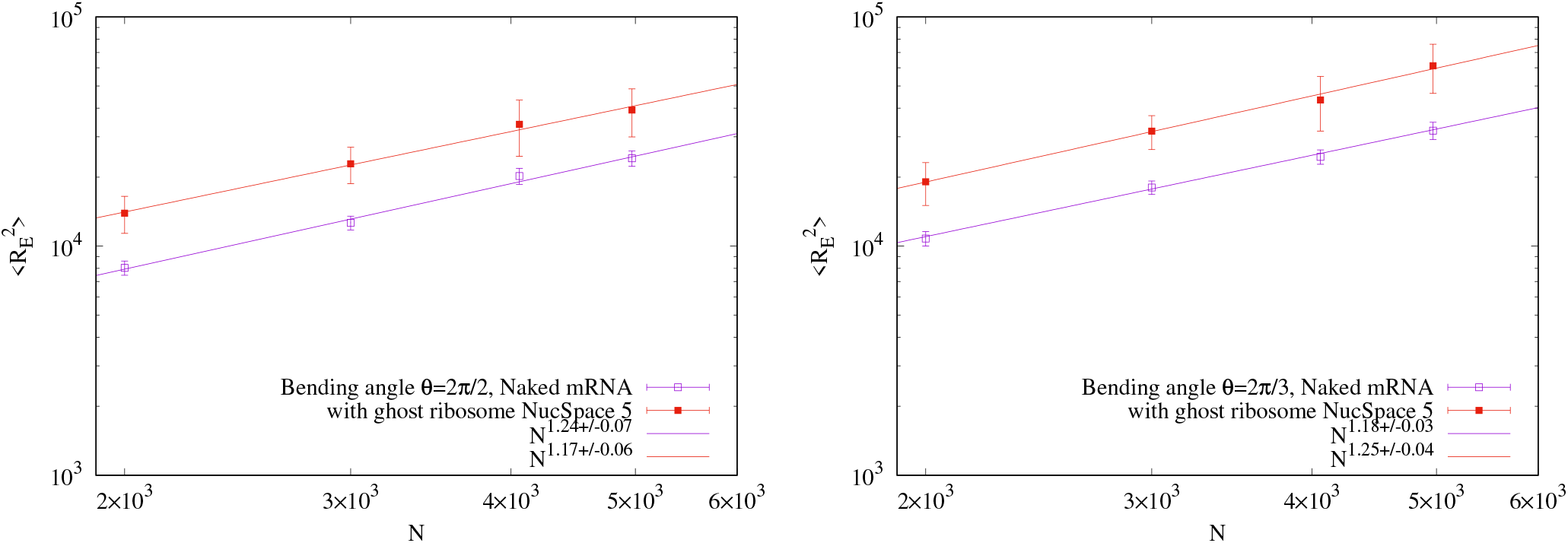
Mean square end-to-end distance 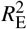 as a function of monomer number *N* for no-ribosome (open square) and ghost-ribosome with *N*_space_ = 5 (close square). The error bars indicate the standard error. Solid and dashed lines represent least-squares fits to *n*^2*v*^. Left: The bending angle *θ* = *π*/2. Right: *θ* = 2*π*/3

In contrast to the “ghost” ribosome cases, the mRNA associated with “real” ribosomes—incorporating repulsive Gaussian core interactions—exhibits a scaling exponent *v* that is larger than the SAW value within the investigated range of *N*, as shown in Fig. 4. While standard polymer theory for dilute solutions predicts that bulky repeating units would eventually act as renormalised beads preserving the SAW universality (*v* ≈0.59), our results show that the explicit excluded volume between ribosomes significantly modifies the backbone statistics in these transcript-scale systems, leading to a more extended conformation.

**FIG. 4.**
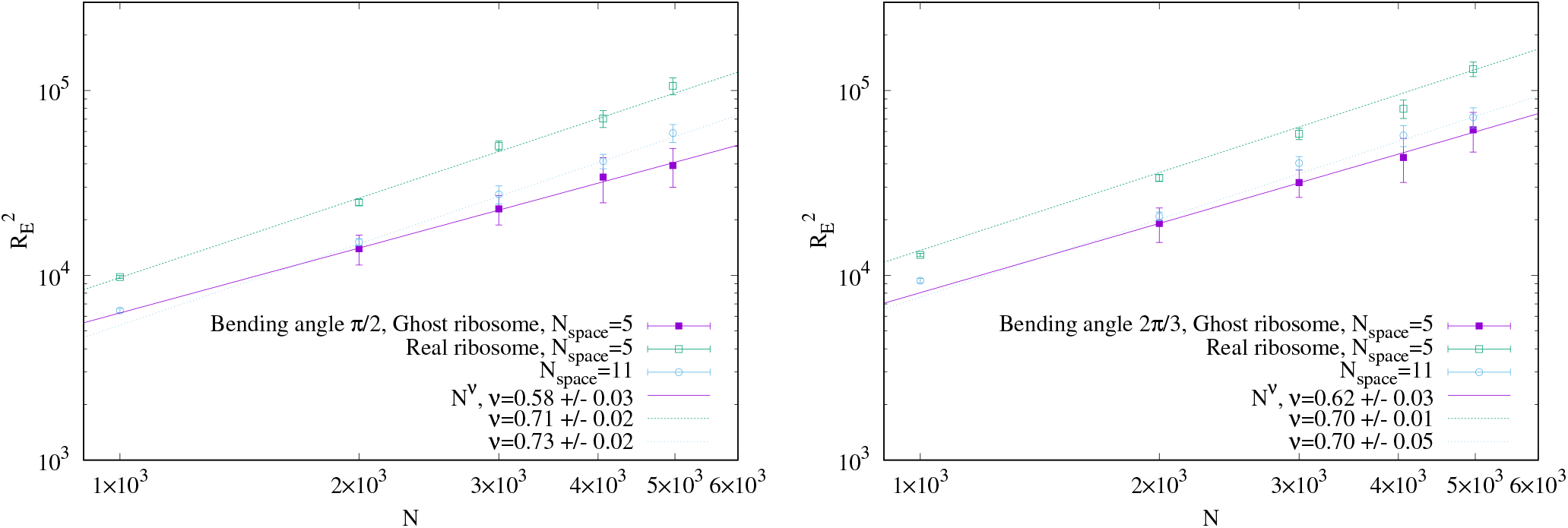
Mean square end-to-end distance 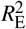 as a function of monomer number *N* for ghost ribosome (closed squares), real ribosome with *N*_space_ = 5 (open squares) and *N*_space_ = 11 (open circle). The error bars indicate the standard error. Solid, dashed and dotted lines represent least-squares fits to *N*^2*v*^. Left: The bending angle *θ* = *π*/2. Right: *θ* = 2*π*/3

To ensure a consistent estimation of the scaling exponent across different densities, we focused on the regime where the number of ribosomes, *M*, is sufficient to establish a collective steric environment. For the lower density case (*N*_space_ = 11) at *N* = 1, 000, the system contains only *M* ≈3 ribosomes. In this small-*M* limit, 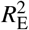 is observed to be slightly larger than the values predicted by the scaling fit for longer chains. This behaviour is distinct from standard homopolymers, where small-*N* chains often appear more compact than the SAW limit. In our model, the massive excluded volume of the ribosomes dominates the local chain statistics; the terminal ribosomes experience an unbalanced outward “push” from the central ribosome without any compensating pressure from neighboring units. This allows the ends to effectively flare outward, leading to a more extended configuration before the collective “steric corridor” of a long polysome is fully developed. Consequently, we exclude this data point from the fitting to focus on the structural properties of polysome architecture with relevant lengths.

For *θ* = *π*/2, the observed value of *v*≈ 0.7 represents a “stretched” conformational state. Within the framework of a Flory-type relation *ν* = 3/(*d*_eff_ + 2), this exponent can be formally associated with an effective dimensionality *d*_eff_ ≈ This suggests that the presence of massive ribosomal volumes restricts the available conformational space for the mRNA backbone, effectively confining the chain within the simulated volume. Such self-induced confinement prevents the backbone from adopting a typical random-coil configuration, resulting in the observed non-SAW statistics.

This sterically-driven expansion is consistently observed across different insertion geometries in our model. For *θ* = 2*π*/3, the scaling exponent remains *v* = 0.70± 0.01 for *N*_space_ = 5 and 0.70± 0.05 for *N*_space_ = 11. The agreement of *v* across different densities and insertion angles in these simulations suggests that the excluded volume of the translation machinery is a primary factor governing the spatial organization of mRNA at biologically relevant lengths. Crucially, for *θ* = 2*π*/3 at the *N*_space_ = 11 case, the averaged distance between ribosomes is larger than the interaction range of the Gaussian core potential (36*u*). The mean inter-ribosomal distance is 40.1*u* (based on 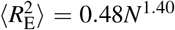 at the sub-chain length *N* = 30*N*_space_ = 330). These results imply that the increase in *v* is not driven by direct ribosome-ribosome repulsion. Instead, it originates from a global change in the effective dimensionality *d*_eff_ imposed by the ribosomal presence along the mRNA strand.

To validate the scaling exponent *ν* through an independent physical property, we calculate the static structure factor *S*(*q*) of the mRNA chain:

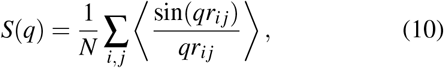

where *q* denotes the magnitude of wave vector and *r*_*i j*_ is the distance between the *i*-th and *j*-th monomers. Note that the coordinates of the ribosomal subunits are excluded from this calculation to focus solely on the statistical properties of the mRNA backbone.

According to polymer scaling theory^26,27^, the structure factor follows a power-law decay *S*(*q*) ∝ *q*^−1/*v*^ in the fractal regime, defined by the conditions *qR*_*g*_≫ 1. This relationship allows for a direct comparison between the scaling xponents *v* obtained from the real-space distance 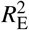 and the reciprocal-space structure factor *S*(*q*).

First, we examine *S*(*q*) for the ghost ribosome system with *θ* = *π*/2, *N*_space_ = 5 and *N* = 4969, as shown in Fig. 5. At intermediate wave vectors (0.03 < *q* < 0.3), the emergence of plateaus and subtle fluctuations indicates a localised departure from self-similarity, reflecting the structural correlations within ribosome-constrained segments. Since our primary interest lies in the global statistical properties of the mRNA backbone, we performed a power-law fit, *S*(*q*) = *Aq*^−1/*v*^, within the range 0.4 < *q* < 2. In this interval, the aforementioned local structural correlations become negligible, and *S*(*q*) exhibits consistent fractal scaling over nearly a decade of the wave vector. Furthermore, this range satisfies the condition *qR*_*g*_ **≳** 20 (estimated using 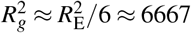).

**FIG. 5.**
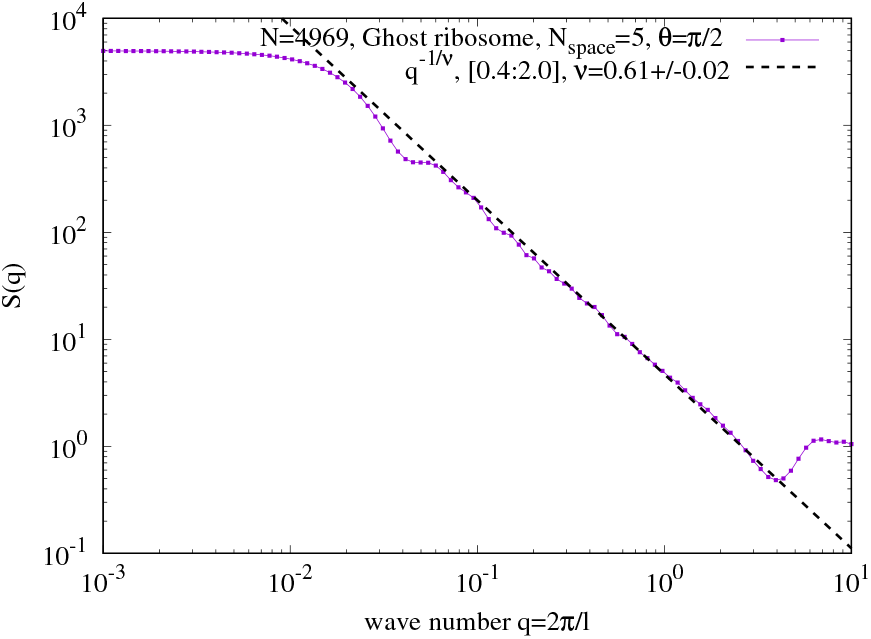
Structure factor *S*(*q*) for ghost ribosomes with *θ* = *π*/2, *N* = 4969, and *N*_space_ = 5. The dashed line shows a least-squares fit to *q*^−1/*v*^, yielding *v* = 0.61 ± 0.02.

The fit yields *ν* = 0.61 ± 0.01, showing good agreement with the data over the broad range of 0.1 < *q* < 3. This result confirms that, in the absence of ribosomal-excluded-volume interactions, the mRNA maintains the SAW universality class. Notably, this exponent is consistent with previous studies of SAW chains on simple cubic lattices with a bending potential *ε*_*b*_(1− cos *θ*) for *ε*_*b*_ < 5^28^.

In the presence of explicit ribosomal excluded volume interactions, the mRNA chain exhibits a robust and consistent scaling behaviour that reflects a significant departure from standard polymer statistics. Figure 6 (Left) shows *S*(*q*) for *θ* = *π*/2 with *N*_space_ = 5 and 11. To ensure rigorous statistical convergence and address potential sampling biases, these results were averaged over 32 independent replicas, with each simulation extending well beyond the decorrelation time of Self-induced Dimensional Reduction and Scaling Transition of mRNA in Polysomes the chain. Remarkably, both *N*_space_ systems display nearly identical behaviour in the fractal regime, demonstrating that the structural expansion of the backbone is a robust consequence of ribosomal crowding, largely independent of the specific ribosome density within the biologically relevant range.

**FIG. 6.**
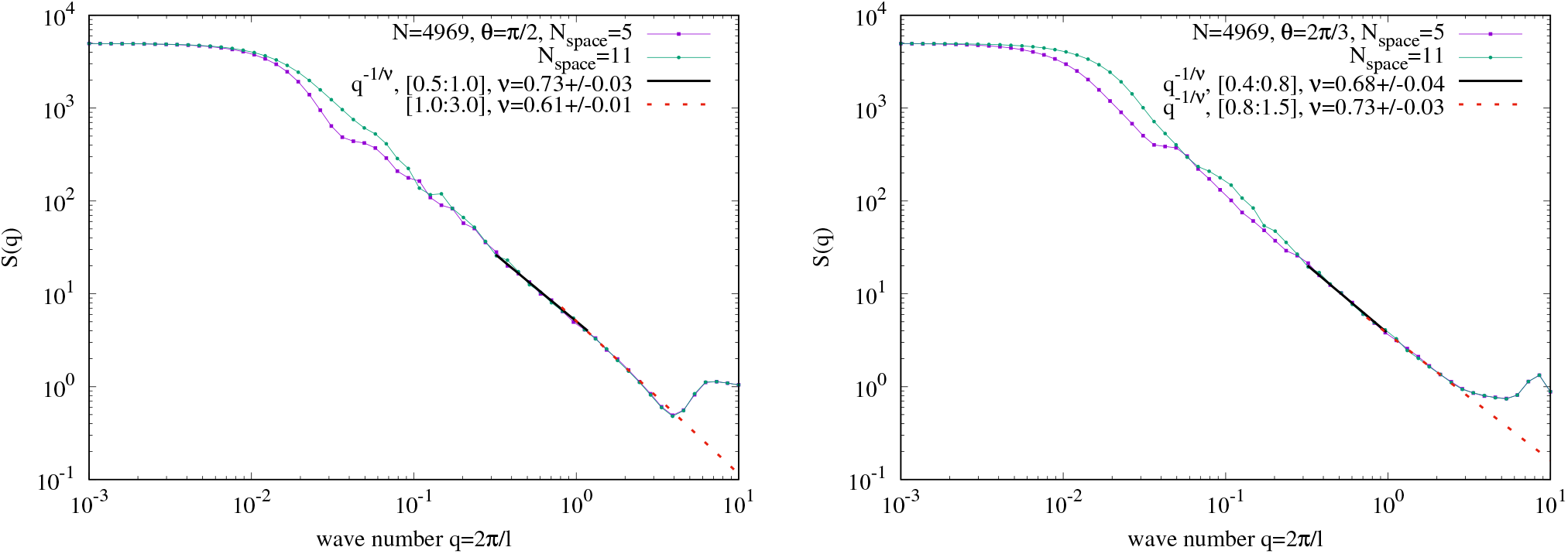
Structure factor *S*(*q*) for real ribosomes with *N* = 4969 and *N*_space_ = 5(square), 11(circle). The solid, dashed and dotted line shows a least-squares fit to *q*^−1/*v*^. Left: The bending angle *θ* = *π*/2. Right: *θ* = 2*π*/3

For the intermediate scale (0.5 < *q* < 1.0), after the transient structural interference from local ribosome insertion and ribosome–ribosome repulsions has subsided, we obtain a scaling exponent of *v* = 0.73± 0.03 for both *N*_space_ cases. This value is significantly higher than both the ghost-ribosome system (*v*≈ 0.61) and the theoretical value for a self-avoiding walk (SAW) in three dimensions (*v* ≈0.59). This provides direct evidence that the ribosomal excluded volume effectively constrains the backbone’s conformational space, forcing it into an inherently more extended, “stretched” state through self-induced confinement.

This local scaling observed in *S*(*q*) is in excellent agreement with our global analysis of the end-to-end distance *R*_*e*_, which yields a consistent apparent exponent of *v*≈ 0.70±0.02 for the range *N* = 1, 000 to 5, 000. The convergence of these two independent measures—local correlation through *S*(*q*) and global extension through *R*_*e*_—strongly supports the emergence of a distinctive conformational regime induced by steric effects.

While the structure factor in the lower-*q* region (*q* < 0.4) exhibits increased statistical noise and is subject to structural interference from the long-range arrangement of ribosomes, the consistency of the power-law decay in the intermediate *q*-range offers a reliable fingerprint of the backbone’s internal architecture. We conclude that while the system may eventually cross over to SAW statistics in the limit of infinitely long chains (*N*→ **∞**), the biologically relevant scale of polysomes is dominated by this self-induced expansion. This mechanism ensures that the mRNA remains in a highly accessible and non-entangled configuration, which is likely essential for efficient translation and protection from enzymatic degradation.

Finally, in the high-*q* regime (1.0 < *q* < 3.0), the exponent reverts to *ν* = 0.61± 0.01, consistent with the universal SAW behaviour. This separation of scales clearly demonstrates that, while the ribosomal network globally stretches the mRNA strand through both geometric and statistical constraints, the intrinsic polymer statistics are preserved at the sub-chain level.

Figure 6 (Right) shows S(q) for the system with a more restricted bending angle, *θ* = 2π/3, at *N*_space_ = 5 and 11. Similar to the *θ* = π/2 case, both systems exhibit almost identical behaviour in the fractal regime regardless of *N*_space_.

For *θ* = 2*π*/3, in the intermediate scale (0.4 < *q* < 0.8) where ribosomal interference is negligible, we obtain a scaling exponent of *v* = 0.68± 0.04 for both *N*_space_ cases. This value is not only significantly higher than that of the ghost-ribosome system (*ν* = 0.61± 0.01) but also the same as the value for the *θ* = *π*/2 system (*v* = 0.73± 0.03) within error. Surprisingly, even when fitting in the higher-*q* range (0.8 < *q* < 1.5), the exponent remains high at *v*≈ 0.6. This persistent scaling indicates that the mRNA is no longer a standard SAW chain and instead behaves as a stretched chain, suggesting a fundamental change in the intrinsic polymer statistics at this angle. We consider that this exponent change is driven by the interplay between the geometric constraints imposed by the bending angles and the ribosomes’ spatial occupancy. The restrictive bending angle confines the bond directions within a narrower cone, which further enhances the reduction of the effective dimension *d*_eff_.

To confirm the enhanced confinement of bond orientations in the presence of real ribosomes, we calculate the bond-bond correlation function *C*(*n*) strictly within the sub-chain segments between two neighbouring ribosomes:

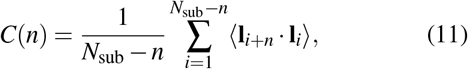

where *N*_sub_ is the number of bonds in a sub-chain (defined as *N*_sub_ = 30(*N*_space_− 1) in our model), and **l**_*i*_ = **r**_*i*+1_ −**r**_*i*_ is the *i*-th bond vector. The summation is restricted to pairs of bonds belonging to the same segment between two consecutive ribosomes.

It is well known that for a simple freely rotating chain (FRC), *C*(*n*) is characterised by a single characteristic length, resulting in a simple exponential decay: *C*(*n*) ∝ exp[*n* log(⟨ cos(*π*− *θ*) ⟩)]. In contrast, our KG model, even in the ghost-ribosome system, exhibits more complex behaviour in the short-range regime due to excluded-volume interactions between monomers, which introduce multiple length scales.

For a swollen polymer chain in the long-range regime, *C*(*n*) is expected to follow the universal scaling relation *C*(*n*) ∝ *n*^2(*v*−1)29–31^. If the real ribosomes effectively restrict the bond orientations, *C*(*n*) is expected to show a significantly slower decay (corresponding to a larger *ν*) compared to the ghost-ribosome system.

Note that, because this analysis is performed within finite-sized sub-chains, a simple power-law decay may not hold over the entire range. As *n* approaches the sub-chain length *N*_sub_, the geometric boundary conditions, where the ends of the chain are tethered to the ribosomes, become dominant. We consider that this confinement will cause the correlation to deviate upward from the power-law decay, eventually exhibiting a “regain” of correlation. Such a departure from standard polymer statistics provides direct evidence that the ribosomal arrangement forces the mRNA strand to transition from a random-walk-like distribution to a more ordered, stretched chain structure within the inter-ribosomal space.

Figure 7(Left) shows *C*(*n*) for *θ* = *π*/2 with *N*_space_ = 5 and 11. In the small-*n* regime (*n* < 10), all systems exhibit strong oscillations. Specifically, *C*(1) ≈ 0.02, consistent with the bending potential, while *C*(2) jumps to approximately 0.3, followed by a gradual decay with oscillations up to *n*≈ 7. This oscillation originates from local excluded-volume correlations and the associated entropic effects. In a configuration where **l**_*i*+2_ = −**l**_*i*_, the possible orientations for the subsequent bond **l**_*i*+3_ are severely restricted to avoid steric overlaps with the preceding segments (notably **l**_*i*_), while still maintaining the bending angle constraint of *π*/2. This significant reduction in the number of accessible microstates leads to a substantial loss of conformational entropy. To avoid such an entropic penalty, the system statistically suppresses the **l**_*i*+2_≈ −**l**_*i*_ state, effectively forcing **l**_*i*+2_ to point away from −**l**_*i*_. This mechanism results in the pronounced positive correlation at *n* = 2 and the characteristic oscillatory profile in the short-range correlation.

**FIG. 7.**
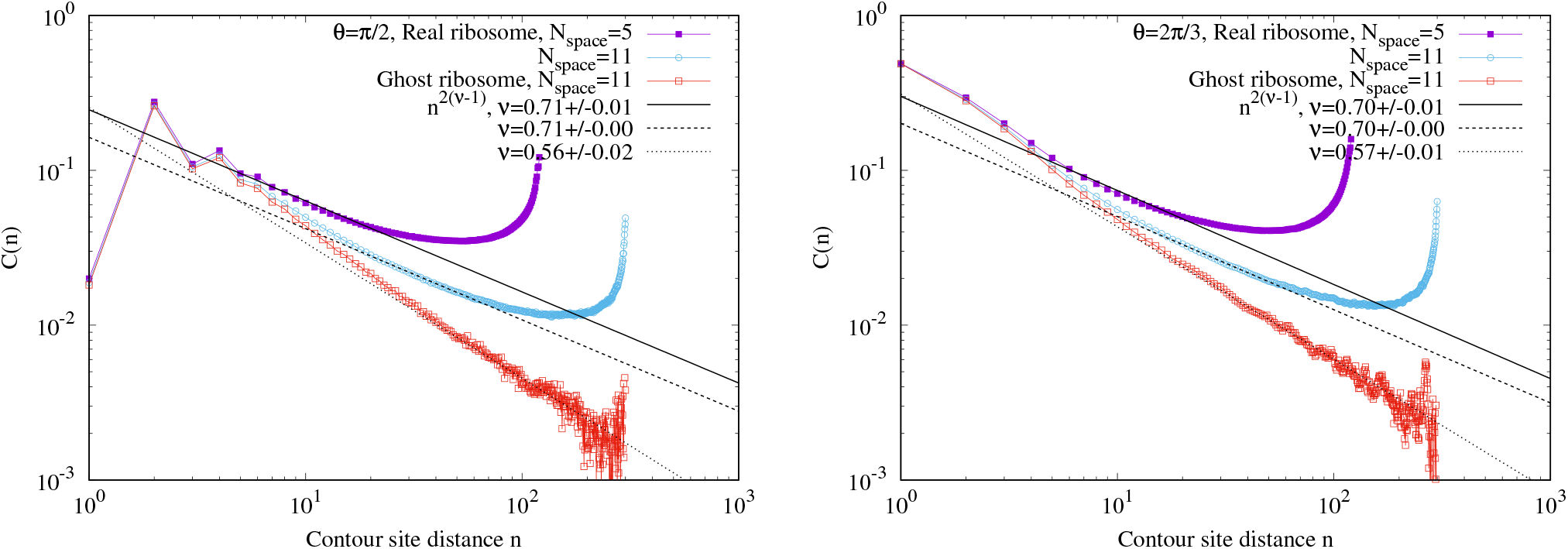
Bond-bond correlation function *C*(*n*) vs. contour site distance *n* for real ribosomes with *N*_space_ = 5 (closed squares) and 11 (open circles), and ghost ribosomes with *N*_space_ = 11 (open squares). Solid, dashed, and dotted lines represent least-squares fits to *n*^2(*v*−1)^. Note that, because this analysis is performed within finite-sized sub-chains, a simple power-law decay may not hold over the entire range. Hence, the obtained exponent is only an effective, rough approximation. Left: *θ* = *π*/2. Right: *θ* = 2*π*/3.

In the large-*n* regime, the behaviour differs markedly between systems. For ghost ribosomes, *C*(*n*) exhibits a monotonic power-law decay with *ν* = 0.56± 0.02, consistent with previous SAW studies. In contrast, real ribosome systems exhibit a plateau followed by a regain of correlation. The plateau begins at *n*≈ 20 for *N*_space_ = 5 and *n* ≈ 100 for *N*_space_ = 11. The regain of correlation is observed from *n* ≈ 100 for *N*_space_ = 5 and *n*≈ 200 for *N*_space_ = 11. Before the plateau, *C*(*n*) follows a power law with *ν* = 0.70 ±0.01, indicating significant stretching.

For *θ* = 2*π*/3, *C*(*n*) exhibits qualitatively similar behavior but with a larger scaling exponent, *v* = 0.70 ± 0.01, in the mid-range (Fig. 7, Right), which is in agreement with the global stretching observed in 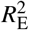 and *S*(*q*).

These results microscopically substantiate the reduction of the effective dimension of the mRNA. The slower decay and the regain of correlation demonstrate that ribosomal excluded volume effectively funnels the chain’s conformational space, forcing it into a stretched chain alignment between ribosomal nodes. In addition, our results clearly demonstrate that the presence of real ribosomes leads to a much slower decay of bond correlations. Moreover, the regain of *C*(*n*) as *n* approaches the boundaries of the sub-chain indicates that the bond orientations are geometrically constrained to align with the axis connecting the ribosomes. This provides direct, microscopic evidence for the reduction of the effective dimension of the mRNA backbone, consistent with the global stretching observed in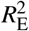 and *S*(*q*).

## IV. DISCUSSION

The striking result of this study is that,in the biologically relevant length scale, the attachment of massive ribosomal constraints fundamentally alters the scaling behaviour of the mRNA backbone, driving the scaling exponent *v* toward the stretched chain (*v*≈ 0.7). It is important to distinguish this from previous numerical studies^32^, which reported that *ν* can reach values near 0.8 in bead-spring models using long-ranged Gaussian core potentials, where the interaction range is 4 times the bead diameter in the longest case. Despite the apparent similarity in the increase of *ν*, the underlying physics in our system is qualitatively different. In such models, the increase in *v* originates from enhanced internal stiffness due to direct, long-range bead-bead repulsion. In contrast, our model relies on short-ranged monomer interactions (WCA). Even with modest-ranged Gaussian core potentials, at an interaction range of 1.2 times the ribosome diameter, the observed increase in *ν* arises from a global reduction in the effective dimensionality *d*_eff_ imposed by the massive ribosomal volumes. This is microscopically substantiated by the “regain” of bond-bond correlations *C*(*n*) at intervals corresponding to neighbouring ribosomes, which is absent in standard homopolymer models, confirming that the ribosomal arrangement funnels the mRNA into a stretched chain alignment.

Experimental observations have established that ribosome or protein binding induces RNA stiffening and structural stabilisation^33,34^. While it may seem intuitive that attaching ribosomes extends the RNA chain by increasing its effective persistence length, our results reveal a more profound mechanism. For instance, numerical work by Fernandes et al.^14^, which examines mRNA recycling rates, assumes that ribosomal attachment enhances the mRNA’s persistence length. This assumption is physically equivalent to our “ghost-ribosome” system, in which appendages only provide local rigidity without mutual interference. From our findings, we already know that, in a system with real ribosomes, the expansion mechanism is fundamentally different. The observed significant extension is not driven by a simple increase of persistence length, but by the reduction of *d*_eff_ due to multi-body excluded volume effects.

Regarding the scaling behavior, we acknowledge that systems with short-range repulsions, such as the Gaussian-core-like potentials used here, are theoretically expected to asymptotically converge to self-avoiding walk (SAW) statistics as *N* → **∞**^35^. True long-range orientational order and non-SAW exponents typically require long-range interactions *U* (*r*) **∼***r*^−*γ*^ with *γ* < 3^35^. However, within the biologically relevant range of *N* investigated in this study, the observed scaling (*v*≈ 0.7) likely represents a robust crossover or pre-asymptotic regime. While this behavior may be formally interpreted as a finite-size effect, it constitutes the dominant physical environment for actual polysomes. Therefore, the steric organization and orientational order reported here are functionally significant features of mRNA at its natural scale, regardless of the system’s hypothetical behavior in the infinite-chain limit. The stiffening mechanism tackled in this work must be distinguished from the classical confinement effects reported in the literature. Standard scaling theories for polymers in confined geometries, such as nanoslits or cylindrical pores^36–38^, rely on *static, external boundary conditions* in which a rigid wall restricts the conformational space. In contrast, the dimensional reduction in our polysome model is *self-induced and internal*. Here, the confinement is dynamically generated by the excluded volume of the ribosomes, which are bulky appendages tethering themselves to the mRNA chain. Notably, our system achieves a nearly 2D limit (*v* ≈0.7) without any physical walls. This suggests that the extreme spatial asymmetry creates an “steric corridor” along the chain’s backbone, representing a distinct class of polymer statistics in which the “wall” and the “chain” form a single dynamical entity.

Although our current mRNA-Ribosome model can not reproduce some complex helical or circular configurations observed in cryo-ET studies^9^, we are working on extending our model for circular configurations. The formation of higher-order architectures requires anisotropic interactions between ribosomal subunits or specific constraints imposed by the exit tunnel. Nevertheless, this work demonstrates that the geometric stretching of mRNA is a robust physical prerequisite for any such ordering. By showing that mRNA stiffening is an unavoidable consequence of ribosomal crowding, this study bridges the gap between universal polymer scaling and the structure of mRNA-Ribosomes machinery during translation. This multiscale phenomenon, made accessible by the tree-based neighbour list algorithm, underscores the importance of accounting for large-scale geometric constraints in the study of cellular supramolecular assemblies.

## V. CONCLUSION

In this study, we have combined a high-performance tree-based neighbour list algorithm with large-scale coarse-grained molecular dynamics simulations to elucidate the conformational statistics of mRNA within a polysome. By modeling the system with a realistic degree of spatial asymmetry (*N* = 4, 969 for the backbone and ribosomes of physiologically relevant volumes), we identified a distinctive expansion in the polymer’s scaling behavior (*v*≈ 0.7) that had previously been overlooked due to modeling and computational limitations. While this observed behavior may represent a crossover or finite-size effect rather than a new universal scaling class in the asymptotic limit, it nonetheless constitutes a critical finding for systems of biologically relevant scales. In the regime where the disparate sizes of the mRNA and ribosomes govern the spatial organization, this structural expansion is the dominant physical reality. Our results provide a fundamental starting point for interpreting recent cryo-ET imaging of polysomes across different cellular environments^8,39^, offering a physical basis for how crowded ribosomal assemblies dictate the three-dimensional architecture of protein synthesis machinery.

Our findings lead to the following key physical insights: (i) the attachment of ribosomes drives the mRNA backbone into a stretched chain regime, characterised by a scaling exponent of *v*≈ 0.7. This exceeds the predictions of previous average-field or “ghost-ribosome” models, which considered only local increases in persistence length; (ii) the stiffening is not an intrinsic property of the chain but is a *self-induced dimensional reduction*. The multi-body excluded-volume interactions between adjacent, bulky ribosomes effectively funnel the mRNA into a quasi two-dimensional “steric corridor” mimicking the effects of external nanoscale confinement without static walls; (iii) the unique periodic “regain” of the bond-bond correlation function *C*(*n*) serves as a structural fingerprint of this transition. It proves that ribosomes act as geometric orientation guides, resetting the chain’s alignment and maintaining long-range order. While our minimal mRNA-Ribosome model focuses on equilibrium excluded-volume effects, it provides a necessary physical foundation for understanding the complex architectures of polysomes. The discovery that ribosomal crowding itself is sufficient to induce a nearly one-dimensional mRNA structure offers a new perspective on how genetic information is physically organised for efficient translation.

Finally, the success of the tree-based neighbour list in handling extreme spatial/size asymmetries opens new avenues for simulating other crowded supramolecular assemblies, such as protein-DNA or protein-RNA complexes. Our work demonstrates that, at the cellular level, large-scale geometric constraints could supersede local chemical details, providing a novel interpretation of polysome organisation.

## ACKNOWLEDGMENTS

We thank Tatsuhisa Tsuboi for illuminating discussions on the experimental measurements of mRNA during translation. H.K. thanks Gemini 3 for its assistance with grammatical correction and prose flow, both of which were subsequently reviewed and approved by the authors. H.V.G. acknowledges financial support from the Ramón y Cajal grant No. RYC2022-038082-I and Spanish Ministry of Science and Innovation, through project PID2023-150536NA-I00, and the “Severo Ochoa” Grant No. CEX2023-001263-S for Centers of Excellence; and CSIC’s grant MMT24-ICMAB-01 for the nanoML4Med project. H.V.G. acknowledges also Red Española de Supercomputación (RES) for the computing time and technical support at the Finisterrae III supercomputer projects FI-2025-2-0058 and Marenostrum 5 FI-2025-3-0049. The authors are also deeply grateful to Kurt Kremer for years of fruitful collaborations and scientific discussions, in particular on the topic of developing efficient methods for multiscale simulations and using polymer concepts in biological systems.

